# Impact of SNP calling quality on the detection of transmission ratio distortion in goats

**DOI:** 10.1101/2021.06.09.447792

**Authors:** María Gracia Luigi-Sierra, Joaquim Casellas, Amparo Martínez, Juan Vicente Delgado, Javier Fernández Álvarez, Francesc Xavier Such, Jordi Jordana, Marcel Amills

## Abstract

Transmission ratio distortion (TRD) is the preferential transmission of one specific allele to offspring at the expense of the other one. The existence of TRD is mostly explained by the segregation of genetic variants with deleterious effects on the developmental processes that go from the formation of gametes to fecundation and birth. A few years ago, a statistical methodology was implemented in order to detect TRD signals on a genome-wide scale as a first step to uncover the biological basis of TRD and reproductive success in domestic species. In the current work, we have analyzed the impact of SNP calling quality on the detection of TRD signals in a population of Murciano-Granadina goats. Seventeen bucks and their offspring (N=288) were typed with the Goat SNP50 BeadChip, while the genotypes of the dams were lacking. Performance of a genome-wide scan revealed the existence of 36 SNPs showing significant evidence of TRD. When we calculated GenTrain scores for each one of the SNPs, we observed that 25 SNPs showed scores below 0.8. The allele frequencies of these SNPs in the offspring were not correlated with the allele frequencies estimated in the dams with statistical methods, thus evidencing that flawed SNP calling quality might lead to the detection of spurious TRD signals. We conclude that, when performing TRD scans, the GenTrain scores of markers should be taken into account to discriminate SNPs that are truly under TRD from those yielding spurious signals due to technical problems.

## INTRODUCTION

In diploid organisms, allelic transmission from parents to offspring is expected to follow the Mendelian law of inheritance, implying that both paternal and maternal alleles are transmitted to the progeny following approximately a 1:1 ratio (Huang et al., 2013). A deviation from this ratio, the so-called transmission ratio distortion (TRD), is produced when either the paternal or the maternal allele is preferentially transmitted to the offspring (Fishman and Mcintosh, 2019). A few years ago, Casellas et al. (2014) implemented a new Bayesian methodology to scan TRD for biallelic markers in diploid organisms. This method was later refined by Vázquez-Gómez et al. (2020) to detect TRD even in pedigrees with incomplete trios. This can be achieved by inferring the probability that specific alleles are present in the parent with a missing genotype based on the allele frequencies of the SNP in the overall population. The accuracy of such inference might be substantially affected by the quality of SNP genotypes. In previous studies (Abdalla et al., 2020; Casellas et al., 2017, 2020 Gòdia et al., 2020; Lahoucine et al., 2020; Vázquez-Gómez et al., 2020), the filtering of SNPs used in TRD scans relied fundamentally on two parameters (genotype call rate and minimum allele frequency or MAF). Deviation from HWE has never been used for this purpose because SNPs showing TRD are expected to display significant departures from HWE. However, it should be noticed that SNPs displaying significant HWE deviations often do so because of genotyping errors (Hosking et al., 2004). In other words, HWE filtering eliminates many unreliable markers which might yield spurious TRD signals just because of technical problems.

The goal of the current work was to make a TRD scan with incomplete trios and then to assess the calling quality of the SNPs that are putatively under TRD by using the GenTrain score implemented in the GenomeStudio software from Illumina (Zhao et al., 2018). The GenTrain score is a metric that fluctuates between 0 (very poor calling quality) and 1 (excellent calling quality) and indicates the reliability of SNP detection based on the distribution of genotypic classes (Pavy et al., 2008; Zhao et al., 2018). By doing so, we aimed to assess the usefulness of the GenTrain score as a complementary metric to be considered in TRD scans as well as to recommend a specific GenTrain score filtering threshold to researchers interested in the detection of TRD.

## MATERIALS AND METHODS

### Sampling and genotyping of Murciano-Granadina goats

As animal material, we have collected blood samples from 17 bucks and their offspring (N=288) in vacuum tubes with K_3_EDTA. These samples have been subsequently stored at −20°C. Since blood collection is a routine procedure performed by CAPRIGRAN, no approval by the Ethics Committee on Animal and Human Experimentation of the Universitat Autònoma de Barcelona was required to perform this experiment. Information about the number of offspring per sire is depicted in Table S1. Genomic DNA extractions were performed following the modified salting out procedure described by Guan et al. (2020). Animals were genotyped with the Illumina Goat SNP50 BeadChip (Illumina Inc., San Diego, CA), which contains 54,241 SNP, following the instructions of the manufacturer. Genotypic data were updated with PLINK 1.9 (Chang et al., 2015) based on the *Capra hircus* genome ARS1 assembly (Bickhart et al., 2017) and the annotation provided by the International Goat Genome Consortium (http://www.goatgenome.org/projects.html#50K_snp_chip). Genotypes were pruned using PLINK 1.9 (Chang et al., 2015). We selected SNPs fulfilling the following criteria: (1) genotype call rate over 95%, (2) minor allele frequency above 0.05 and (3) no missing genotypes in any of the 17 sires. Besides, the percentage of sires heterozygous for each SNP was estimated from the output obtained with the --hwe command of PLINK 1.9 (Chang et al., 2015) in order to remove SNPs with less than 20% of heterozygosity in the sire population (only SNPs with heterozygous genotypes are informative). The reference allele in this subset of the population was set as the most common allele in all individuals.

### Statistical methods

To estimate TRD, we used a frequentist modification (Vázquez-Gómez et al., 2020) of the Bayesian method implemented by Casellas et al. (2014). Assuming two alleles (A1 and A2) and the existence of genotyped heterozygous sires and of ungenotyped dams, this method allows to compute for every marker an α-value which ranges between −0.5 (the A1 allele is not transmitted) and 0.5 (the A2 allele is not transmitted) thus providing an estimate of the magnitude of TRD. Allele frequencies in the ungenotyped dams were inferred by calculating a π-parameter which varies from 0 to 1. The two α and π parameters were estimated by maximizing the likelihood function and the statistical significance of α was assessed by using a likelihood ratio test (Nelson, 2008). A correction for multiple testing was applied to the *P*-values obtained from the χ^2^ distribution using the false discovery rate approach (FDR) reported by Benjamini & Hochberg (1995) to obtain the corresponding *q-values*. Markers with α-values above 0.15 or below −0.15 and *q-values* < 0.05 were considered to show significant TRD. After quality control based on genotype call rate and MAF, the final genotypic data included 42,272 autosomal SNPs.

## RESULTS AND DISCUSSION

The implementation of the TRD test allowed us to identify 2,944 SNPs that were deviating from the Mendelian ratio (α-value > 0.15 or < −0.15). The highest α-value detected in our population was 0.499, while the lowest was −0.428; but for the majority of the genotyped SNPs, α-values were comprised in the [−0.15 to 0.15] interval which indicates the absence of TRD (Fig. 1a). After applying a likelihood ratio test and FDR correction, 36 SNPs were selected as significant (*q-value* < 0.05), from which 15 SNPs had an α-value below −0.15 implying a major transmission of the alternative allele, while the remaining 21 SNPs showed an overtransmission (α-value over 0.15) of the reference allele (Fig. 1b).

**Fig. 1a.**
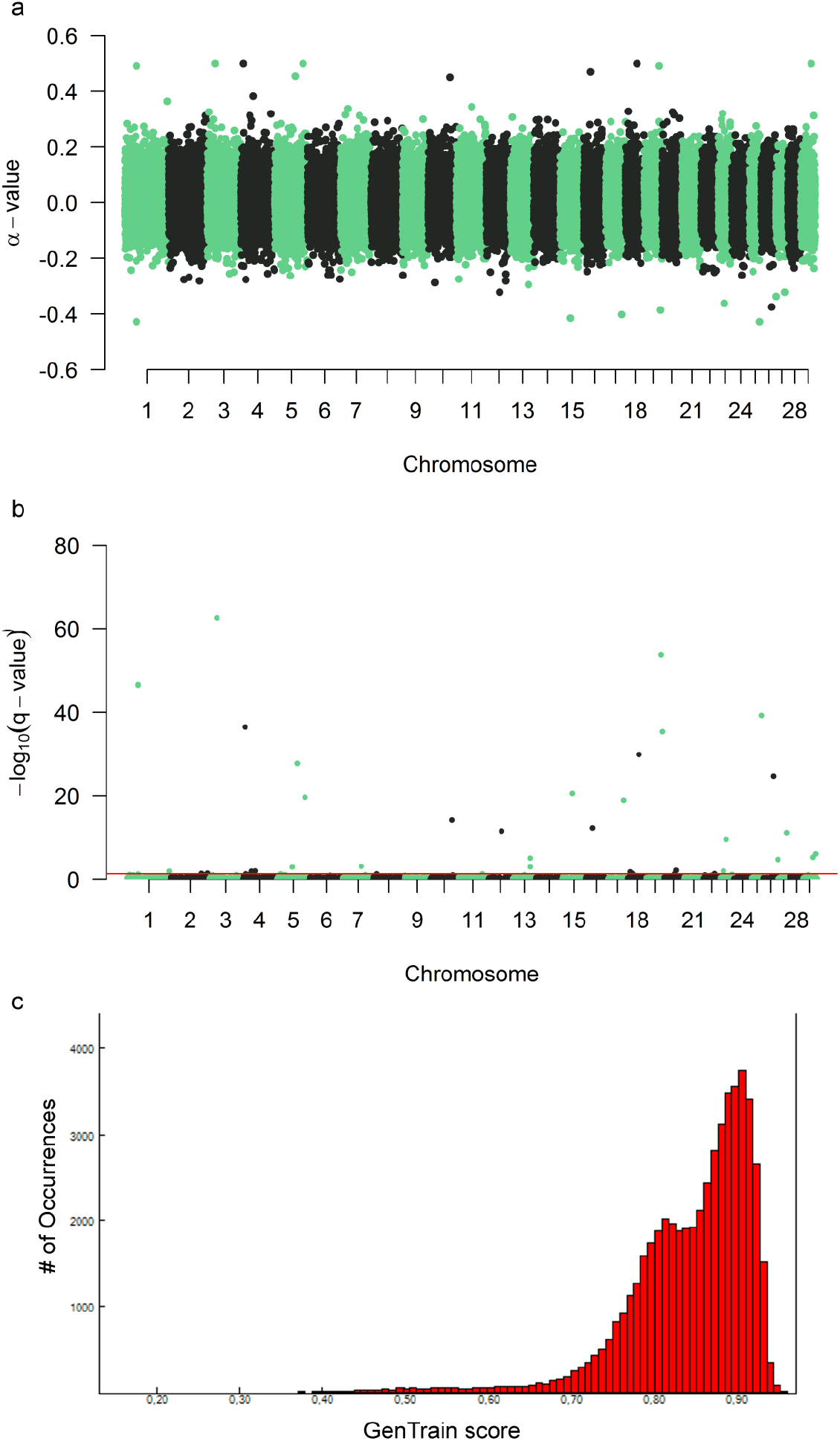
Genome-wide detection of SNP markers that show evidence of transmission ratio distortion in a population comprising 17 sire-families of Murciano-Granadina goats. The α-value estimated for each SNP is plotted in the *y*-axis while the chromosomal locations of SNPs are indicated in the *x*-axis. **1b.** Manhattan plot indicating the statistical significance (y-axis), expressed as −log_10_ of the *q*-value, of the α-values calculated for 42,272 SNPs genotyped in a population comprising 17 sire-families of Murciano-Granadina goats. The chromosomal location of each SNP is indicated in the x-axis. The red line corresponds to the threshold of significance which corresponds to a *q*-value = 0.05 expressed in a −log_10_ scale. **1c.** GenTrain score distribution for 42,272 SNPs genotyped in 305 goats. It can be seen that the vast majority of SNPs have GenTrain scores above 0.70.

In order to verify the accuracy of the genotyping of the 36 SNPs displaying significant TRD, we calculated their GenTrain scores with the GenomeStudio software (Illumina Inc., San Diego, CA). Clustering of SNPs with different GenTrain scores and distribution of the GenTrain scores of the genotyped SNPs are depicted in Fig. 1c and Fig. S1. In Table 1, it can be seen that 25 of these SNPs have GenTrain scores below 0.80, with values ranging from 0.16-0.63 and an average score of 0.51 ± 1.14 (**Group 1**). In contrast, eleven SNPs have GenTrain scores above such threshold, with an average value of 0.87 ± 0.04 (**Group 2**). For each of these two groups of SNPs, we have calculated the correlation between allele frequencies of the SNP in the offspring vs allele frequencies inferred for the ungenotyped dams with the methods reported by Vázquez-Gómez et al. (2020). In principle, allele frequencies of parents and their offspring should be significantly and positively correlated. In the **Group 1** of SNPs such correlation was very weak and non-significant (r = −0.007, *P*-value = 0.9733). Even worse, when we retrieved from **Group 1** seventeen SNPs with GenTrain scores between 0.5-0.6, the correlation was −0.0932 (*P*-value 0.7217). In strong contrast, allele frequencies of mothers and offspring were highly correlated in the **Group 2** of SNPs (r = 0.8656, *P*-value = 0.0005). This result implies that the method implemented by Casellas et al. (2014) and subsequently modified by Vázquez-Gómez et al. (2020) works very well in reconstructing allele frequencies in parental individuals without genotypes when SNPs have high GenTrain scores (> 0.80 in our study), which are the vast majority (Fig. 1c).

**Table 1.**
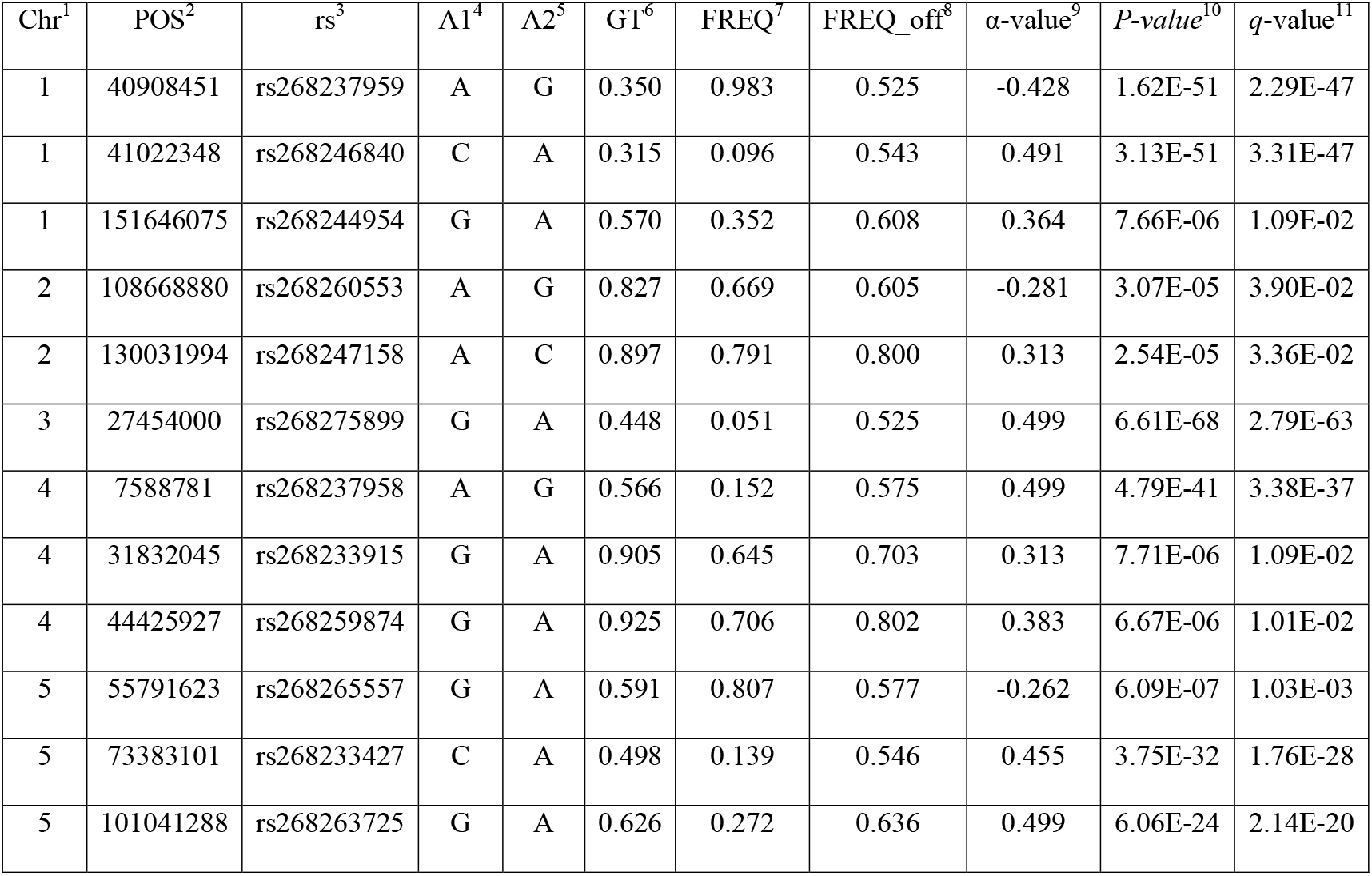
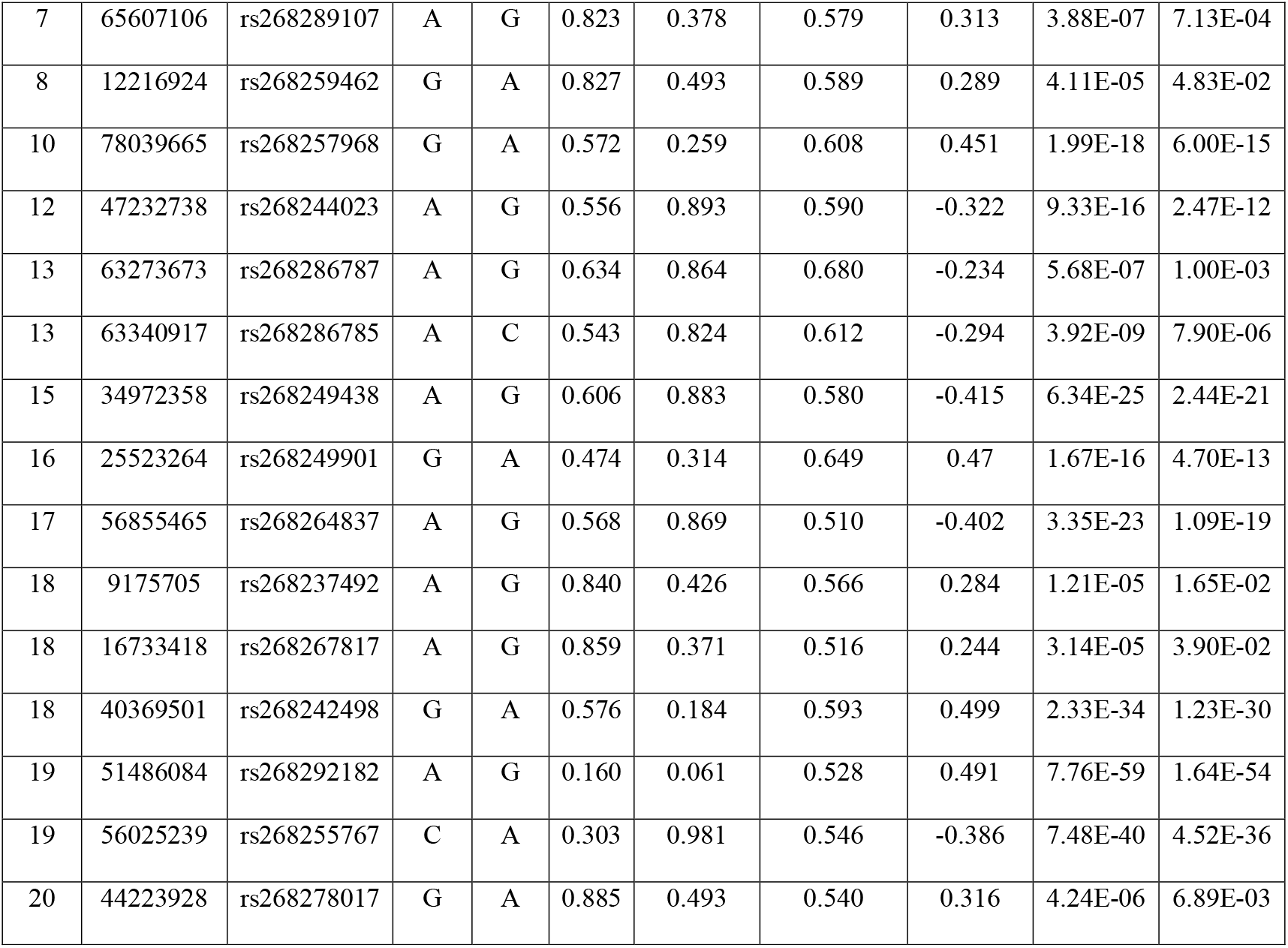
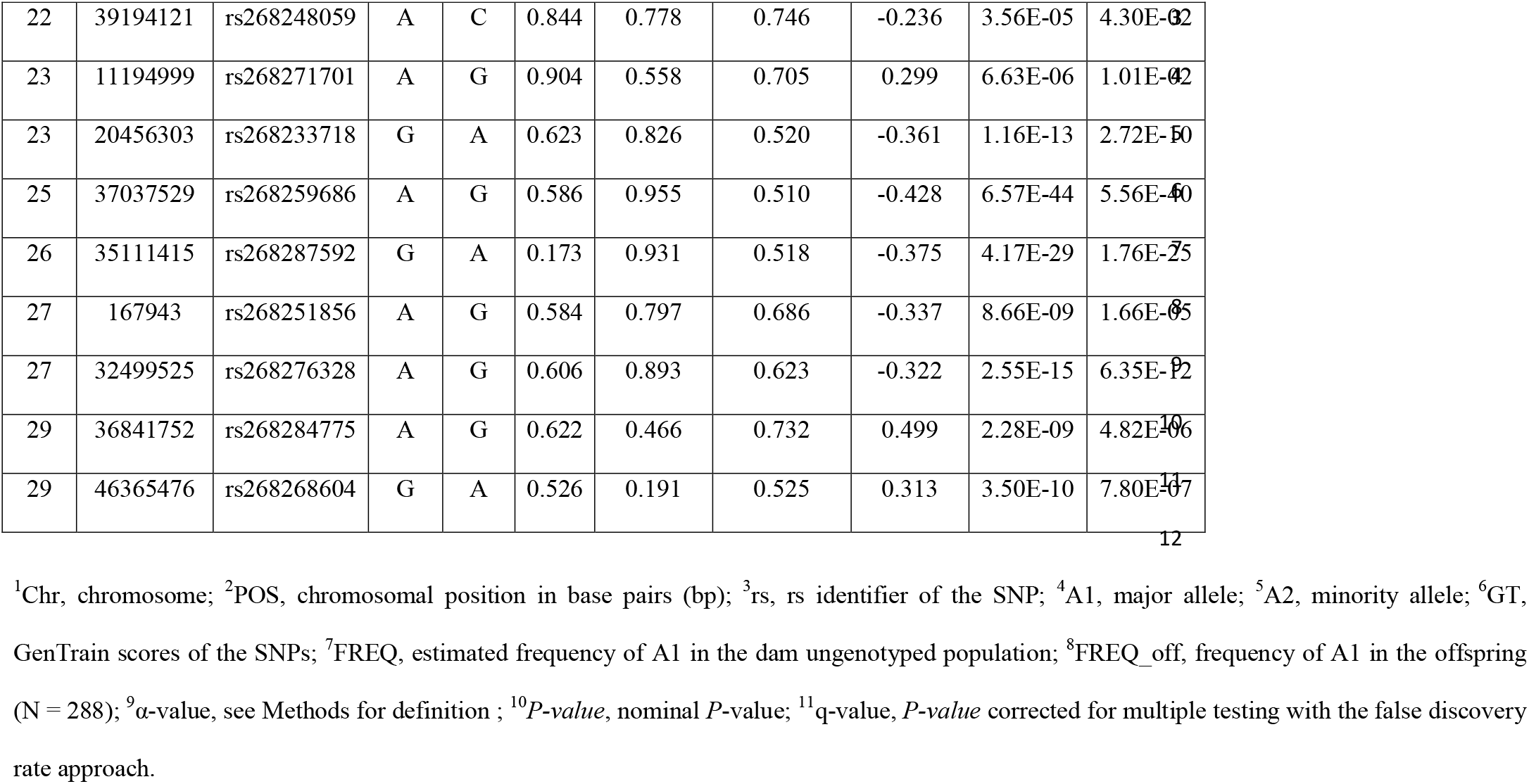
Single nucleotide polymorphisms displaying transmission ratio distortion (α-value above 0.15 or below −0.15, *q*-value < 0.05) in a population composed by 17 families of Murciano-Granadina goats.

Previous reports have indicated that SNPs with minimum GenTrain scores of 0.25 can be safely used in most analyses (Pavy et al., 2008). Indeed, Pavy et al. (2008) designed highly-multiplexed SNP arrays for the genotyping of black and white spruce and reported that SNPs with GenTrain scores of 0.25 are very reliable and have a low rate of missing data. According to Pavy et al. (2008), SNPs with GenTrain scores of 0.25 or more had average call rates above 99%, and for SNPs with GenTrain scores above 0.4 the rate of missing data became negligible and the average rate of missing data per successful SNP was very low, On the other hand, Guo et al. (2014) indicated that SNPs with GenTrain scores above 0.70 are correctly clustered in most cases, while the clustering of SNPs with GenTrain scores below 0.7 might be more problematic. Based on our results, we conclude that the performance of TRD scans, especially in the case in which full trios are not available, should rely on the establishment of a stringent threshold for SNP calling quality. In our study, family size was small (on average 17-18 offspring per sire) and the density of the Goat SNP50 BeadChip (Illumina) is modest. In these conditions, we advise to consider markers with GenTrain scores of 0.80 or higher.

In studies with a more optimal design (particularly regarding family size), such threshold could be determined empirically by analyzing the correlations between the allele frequencies estimated in the parental class without genotypes and those inferred experimentally in the offspring, which should be significant and positive. This simple approach should facilitate the elimination of spurious TRD signals produced by technical factors in order to concentrate efforts on those that have biological implications.

## ACKNOWLEDGMENTS

Many thanks to CAPRIGRAN for carrying out phenotype recording and blood sample collection in Murciano-Granadina goats. This research was funded by the European Regional Development Fund (FEDER)/Ministerio de Ciencia e Innovación - Agencia Estatal de Investigación/Project Reference grant: PID2019-105805RB-I00 and by the CERCA Programme/Generalitat de Catalunya. We also acknowledge the support of the Spanish Ministry of Economy and Competitivity for the Center of Excellence Severo Ochoa 2020-2023 (CEX2019-000902-S) grant awarded to the Centre for Research in Agricultural Genomics (CRAG, Bellaterra, Spain). We also acknowledge the support of the CERCA programme of the Generalitat de Catalunya. Maria Luigi-Sierra was funded with a PhD fellowship “Formación de Personal Investigador” (BES-C-2017-079709) awarded by the Spanish Ministry of Economy and Competitivity.

## AVAILABILITY OF DATA

Genotypes of the 305 Murciano-Granadina sires and offspring and pedigree information are available in 10.6084/m9.figshare.14686230.

