## Supplemental information for "Impact of SNP calling quality on the detection of transmission ratio distortion in goats"

### SUPPORTING INFORMATION

**Table S1.** Family size for each one of the 17 Murciano-Granadina sires.

| ID sire | Number of daughters | Number of sons | Total Offspring |
|---|---|---|---|
| 26137 | 7 | 3 | 10 |
| 47067 | 5 | 5 | 10 |
| 47227 | 5 | 7 | 12 |
| 49946 | 23 | 0 | 23 |
| 49952 | 21 | 0 | 21 |
| 56475 | 12 | 3 | 15 |
| 60401 | 13 | 11 | 24 |
| 148342 | 14 | 0 | 14 |
| 190384 | 24 | 0 | 24 |
| 190580 | 24 | 2 | 26 |
| 203008 | 11 | 0 | 11 |
| 203511 | 8 | 2 | 10 |
| 871651 | 10 | 0 | 10 |
| 871653 | 5 | 6 | 11 |
| 871674 | 27 | 3 | 30 |
| 871680 | 13 | 1 | 14 |
| 871681 | 11 | 12 | 23 |

**Fig. S1.** GenoPlots of SNPs with different GenTrain scores (GT-scores). It can be seen that when the GT-score is high (0.90), the clustering of each of the three genotypes is quite tight. In contrast, SNPs with GT scores of 0.55 and 0.62 display scattered patterns of clustering for at least one of the genotypes. For the SNP with a GT score of 0.17, the pattern of clustering is not credible.

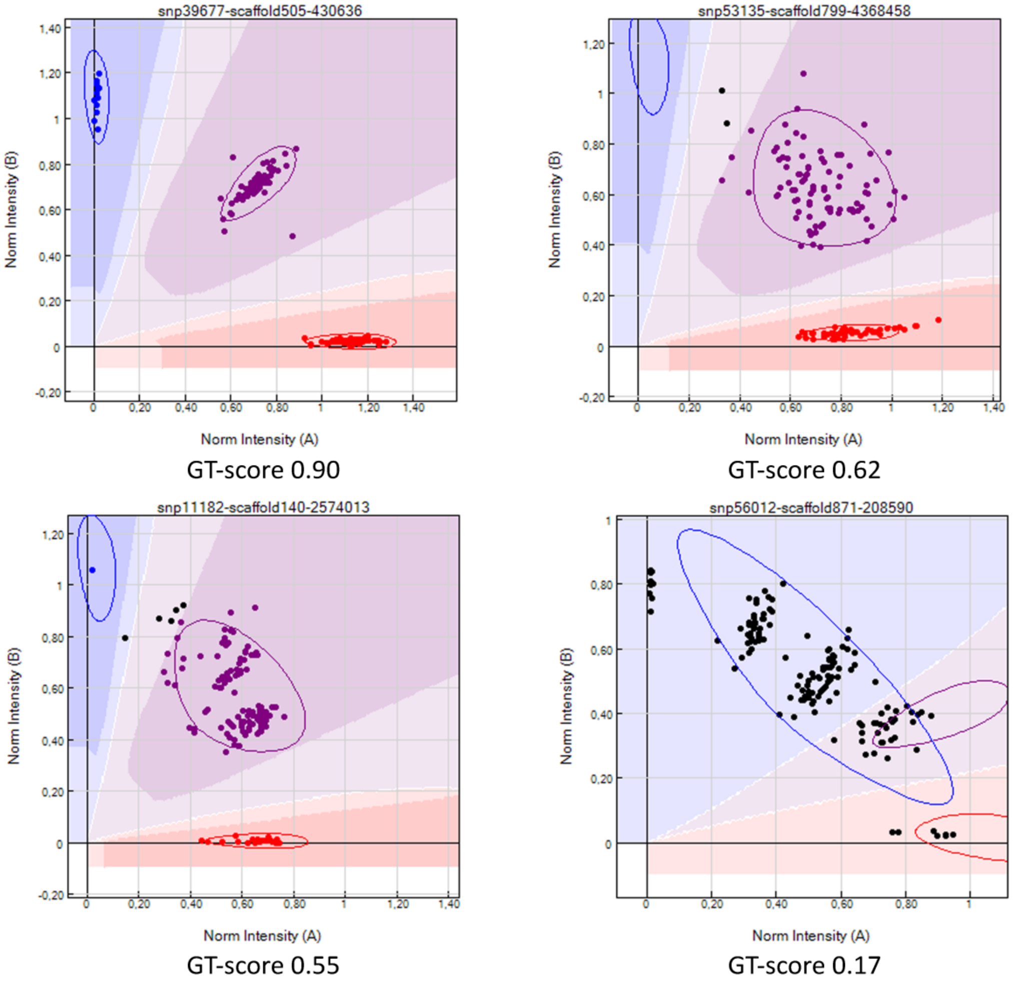
